# Formation of multiple flagella caused by a mutation of the flagellar rotor protein FliM in *Vibrio alginolyticus*

**DOI:** 10.1101/2022.04.21.489128

**Authors:** Michio Homma, Norihiro Takekawa, Kazushi Fujiwara, Yuxi Hao, Yasuhiro Onoue, Seiji Kojima

**Author notes:** Corresponding author: Division of Biological Science, Graduate School of Science, Nagoya University, Chikusa-ku, Nagoya 464-8602, Japan.; E-mail addresses (M.H.) and (S.K.). College of Life Sciences, Department of Bioinformatics, Ritsumeikan University, 1-1-1 Noji-higashi, Kusatsu, Shiga 525-8577, Japan (Yasuhiro Onoue).

## Abstract

The marine bacterium *Vibrio alginolyticus* forms only a single flagellum at the cell pole. In *Vibrio*, two proteins (GTPase FlhF and ATPase FlhG) regulate flagellar number at the cell pole. We previously isolated a mutant strain characterized as NMB155 that forms multiple flagella despite the absence of mutations in *flhF* and *flhG*. NMB155 also exhibited straight swimming without a directional change in flagellar rotation. Whole-genome sequencing of NMB155 identified an E9K mutation in FliM that is a component of the C-ring in the flagellar rotor. Mutations in FliM result in defects in flagellar formation (*fla*) and flagellar rotation (*che* or *mot*); however, there are few reports indicating that FliM mutations increase the number of flagella. Here, we determined that the E9K mutation confers the multi-flagellar phenotype and also the *che* phenotype. The co-expression of wild-type FliM and FliM-E9K indicated that they were competitive in regard to determining the flagellar number. It had been shown that the ATPase activity of FlhG corresponds to the flagellar number. We observed that the ATPase activity of FlhG was increased by the addition of FliM but not by the addition of FliM-E9K. This indicates that the N-terminal region of FliM that includes the E9 residue interacts with FlhG to increase its ATPase activity, and the E9K mutation may inhibit this interaction. We concluded that FliM downregulate FlhG activity to inhibit the formation of additional flagella.

**Importance:** The flagellar rotor generates a driving force to rotate the flagellum and is not involved in controlling the number of flagella in *Vibrio*. However, we observed that the E9K mutation in the rotor protein FliM confers multiple flagella. Our findings reveal a novel regulatory mechanism controlling flagellar number.

## Introduction

Numerous types of bacteria possess motility organs termed flagella on their cell surface. By rotating these flagella in a screw-like motion, the bacteria can swim in liquid or swarm to spread on the surface to move toward favorable or away from unfavorable environments to allow for survival. The flagellum is composed of more than 30 proteins (1). A given flagellum consists of three components that include a motor that is embedded in the membrane, a filament extending outside of the cell, and a hook connecting the motor and the filament. The rotational force of flagellum is generated by converting the influx of coupling ions caused by electrochemical potentials across the cell membrane (2-4). The motor is constructed into two parts that include a stator and a rotor. When ions flow through the stator, the stator undergoes a structural change, and interactions occur between the stator and rotor, ultimately resulting in the rotation of the rotor.

The motor can rotate both counterclockwise (CCW) and clockwise (CW). The direction of rotation is determined by the association and dissociation of the chemotaxis signal protein CheY to and from the motor. The external environment is sensed by chemoreceptors present on the cell membrane, and the signal from the chemoreceptor is transmitted via intracellular proteins through a relay of phosphate groups that includes both phosphorylation and dephosphorylation to CheY (5, 6). Phosphorylated CheY (CheY-P) binds to the motor to cause CW direction rotation, and dephosphorylated CheY moves away from the motor to cause CCW rotation.

The positions and number of flagella vary among bacterial species and are strictly regulated in each species. For example, *Vibrio* spp. and *Pseudomonas aeruginosa* possess one flagellum on one pole of the cell, while *Campylobacter jejuni* possesses one flagellum on both poles and *Helicobacter pylori* possesses multiple flagella on one pole. *Escherichia coli*, *Salmonella*, and *Bacillus subtilis* also possess multiple flagella on their entire cell surfaces (7). Flagellar structure formation begins with the formation of a rotor embedded in the membrane (Fig. S1). Several protein complexes are located in the rotor and extend from the cytoplasm to the outer membrane. Flagellar formation begins with the formation of a complex that is embedded in the inner membrane, the MS-ring, and the export-gate complex. The MS ring is composed of 34 transmembrane protein FliF molecules. In the flagellum of *Salmonella*, it has been demonstrated that overexpression of the FliF protein results in ring formation in *E. coli* cells (8); however, the presence of an export gate complex consisting of core FliP/FliQ/FliR proteins is responsible for efficient ring formation. The C-ring that is constructed at the cytoplasmic interface of the MS-ring plays an important role in the generation of rotational force and control of rotational direction (9-15).

In our laboratory, we investigated flagellar formation in *V. alginolyticus* that possesses only a single flagellum on one pole, and we revealed that FlhF and FlhG are the primary regulators of the number and position of flagella in *V. alginolyticus* (16-18). FlhF and FlhG (also termed FleN or YlxH) are members of the SIMIBI class of nucleotide-binding proteins that include the signal sequence-binding protein Ffh, the signal-recognition particle (SRP) receptor FtsY, and the cell division protein MinD (7, 19). Overexpression of FlhF in *V. alginolyticus* results in an increase in flagellar number, while loss of FlhF results in a lack of flagella. This indicates that FlhF positively regulates the number of flagella (17). Conversely, overexpression of FlhG in *V. alginolyticus* cells results in a lack of flagella, and the loss of FlhG results in an increase in flagellar number, thus indicating that FlhG negatively regulates the number of flagella (17). FlhF is also important in regard to determining the location of flagellar formation, and FlhF molecules themselves may promote flagellar formation at the poles (18, 20). In the absence of FlhF, the polar flagella of *V. alginolyticus* cells are peritrichously generated by the *sflA* mutation (21). The homologous proteins of FlhF and FlhG that are not present in *E. coli* and *Salmonella* but are present in *P. aeruginosa* and *Shewanella putrefaciens* that possess a single polar flagellum exhibit functions that are similar to those of *V. alginolyticus* (22). In *S. putrefaciens,* FlhG interacts with the N-terminal region of FliM, and the N-terminal deletion of FliM confers the multi-flagellar phenotype that is similar to the *flhG* mutant phenotype. The N-terminus of FliM contains a highly conserved motif (EIDAL motif) that is necessary and sufficient for interaction with FlhG (23, 24). FlhG interacts with FlhF in the cytoplasm to inhibit its polar localization. FlhG also interacts with FlhF at the cell pole to inhibit FlhF activity, and this inhibition appears to require the scaffold protein HubP that is localized at the poles (25). Although interactions between FlhG and HubP have been reported, the detailed mechanisms underlying flagellar formation and positioning remains unclear.

We previously isolated a mutant strain exhibiting multiple polar flagella through the use of EMS mutagenesis experiments, and we characterized this strain as NMB155 (Fig. 1) (26). The mutation site for the multi-flagellar phenotype of NMB155 has been a mystery for more than 20 years. Whole-genome sequencing revealed mutations at multiple locations in the chromosomal DNA of NMB155 as previously reported (27). The only mutation involved in flagellar formation that was specifically determined in NMB155 was a point mutation that substituted Glu9 in FliM for Lys (E9K). FliM is a component of the flagellar rotor C-ring. Until now, mutations in FliM have been well established to result in the *fla*, *che*, and/or *mot* phenotypes, and there have been few reports detailing the observation that FliM mutations increase the number of flagella in specific species. Therefore, it is difficult to conclude that the *fliM* gene is responsible for the formation of multiple polar flagella.

**Fig. 1.**
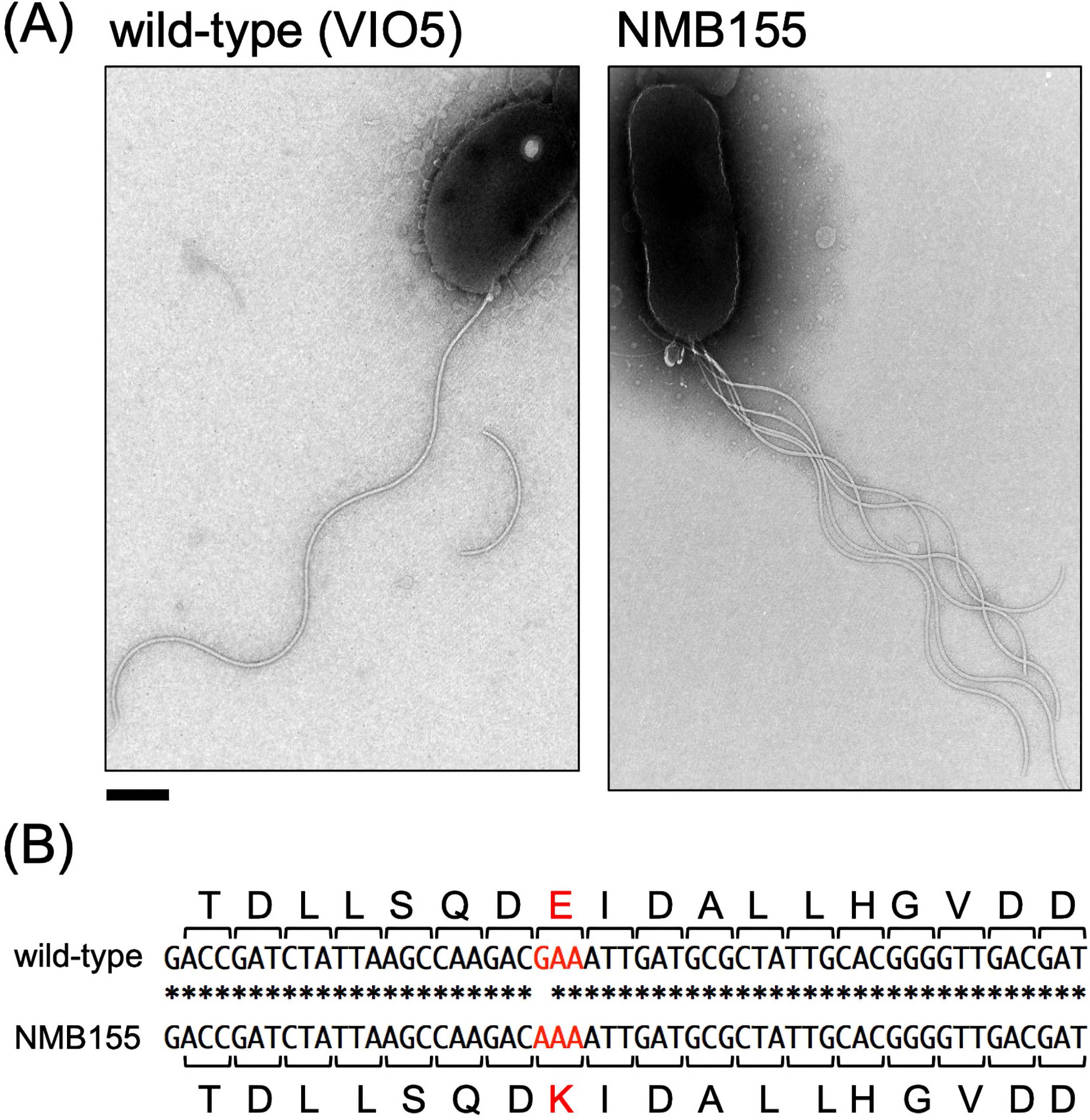
(A) Electron micrographs of the wild-type strain (VIO5) (left panel) and the mutant strain (NMB155) (right panel). Scale bar, 0.5 μm. (B) The mutation site of NMB155. The nucleotide sequences of *fliM* and their amino acid sequences from wild-type (VIO5) and NMB155 are provided.

In this study, we first tested if FliM-E9K is responsible for the multiple polar-flagella phenotype of NMB155, and we aimed to clarify how mutations in FliM, particularly in the E9 site or E9K, are involved in the mechanism of flagellar number control in *V. alginolyticus*.

## Results

### FliM-E9K is the responsible mutation underlying multipolar flagella of NMB155

The entire genome of NMB155 was sequenced as a chemotaxis mutant strain, and an E9K mutation was identified in FliM, a flagella-related gene (27). We amplified the genomic region containing *fliM* of NMB155 using PCR and confirmed the sequence. The *fliM* gene of NMB155 possesses a point mutation in which the 25th guanine (G) from the 5’ side is altered to adenine (A), and as a result, the 9th glutamate (E) of the FliM protein is mutated to lysine (K) (Fig. 1B). The amino acid sequence alignment of the FliM proteins from several bacteria exhibited a motif termed the EIDAL motif (23), and E9 is highly conserved in *Vibrio* and is also in evolutionarily distant species (Fig. S2). To assess if the FliM-E9K mutation was responsible for the phenotype of the multiple polar flagella in NMB155, the *fliM*-deletion mutant strain (Δ*fliM*, NMB321) was transformed with plasmids encoding wild-type *fliM* or *fliM*-E9K. The culture solutions of each strain were spotted on to VPG soft-agar plates, and swimming abilities were measured by observing the spread of colonies (Fig. 2). The sizes of the motility rings in the wild-type strain (VIO5) with all three plasmids were nearly the same. Δ*fliM* with an empty vector did not spread due to its inability to move. Δ*fliM* with the plasmid expressing wild-type FliM spread well but not as much as the wild-type strain. The size of the motility ring of NMB155 possessing the empty vector was similar to that of Δ*fliM* with the plasmid expressing FliM-E9K.

**Fig. 2.**
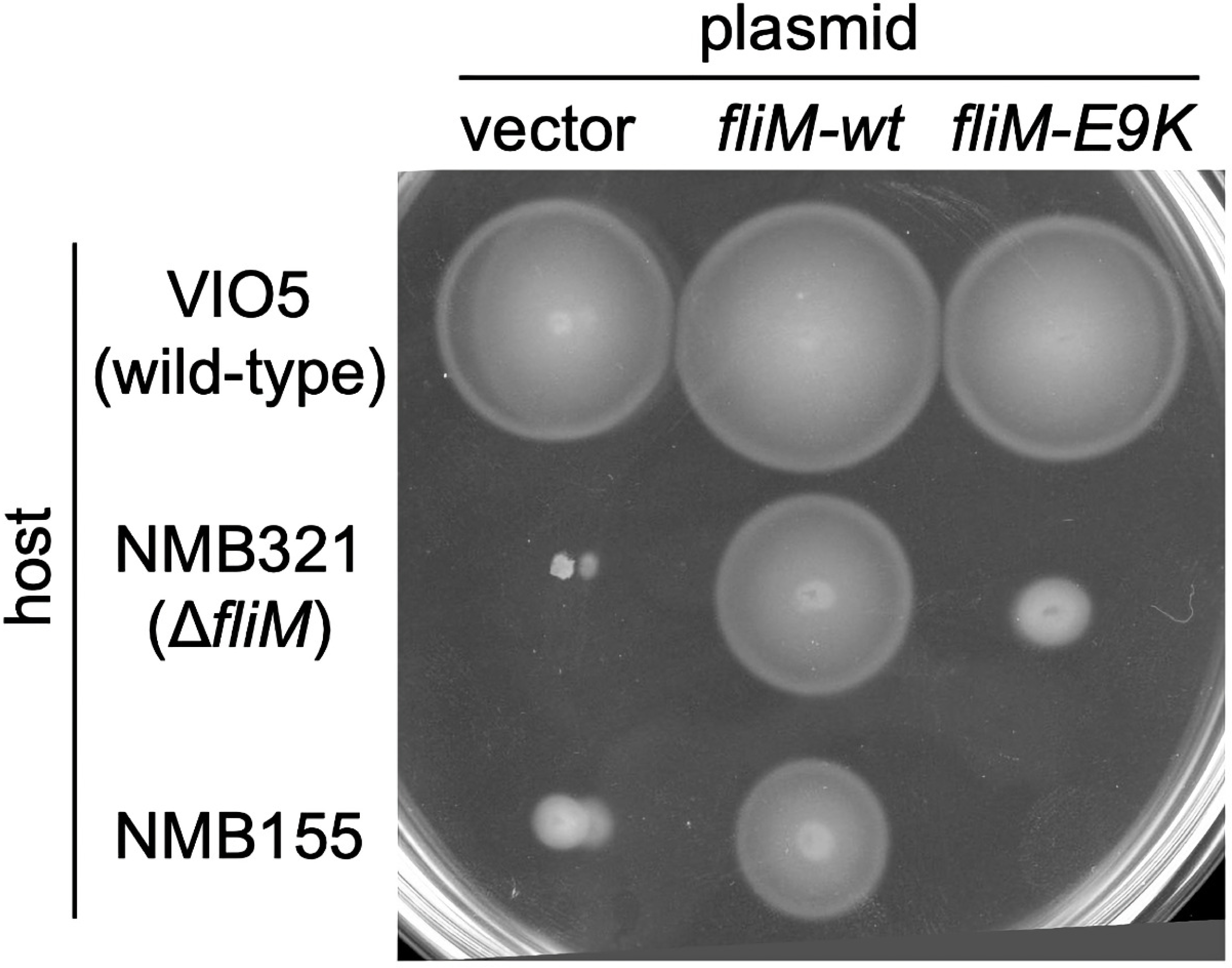
The motility of the FliM-E9K mutant. The fresh colonies of VIO5 (wild-type strain), NMB321 (Δ*fliM* strain), and NMB155 containing pBAD33 (vector), pMK1001 (*fliM-wt*; expressing wild-type FliM), or pMK1001-E9K (*fliM-E9K*; expressing FliM-E9K) were spotted onto VPG soft-agar plates containing 0.02% (w/v) arabinose and 2.5 μg/mL of Cm and then incubated at 30°C for 6 hours.

### Changes in the number of flagella as a result of the FliM-E9K mutation

The number of flagella in each strain was measured using a high-intensity dark field microscope (Fig. 3). The cells of the wild-type strain (VIO5) predominantly possess only one polar flagellum, while more than 70% of NMB155 cells and the *flhG* mutant (KK148) cells exhibit multiple polar flagella; however, the ratio of cells possessing greater than four flagella was higher in the *flhG* mutant than it was in NMB155. Expression of the FliM-E9K protein in Δ*fliM* (NMB321) conferred multiple polar flagella, and this was similar to observations in NMB155. In contrast, the ratio of cells possessing a single polar flagellum was not markedly decreased, and many cells with a single polar flagellum were observed when FliM-E9K was expressed in the wild-type strain. A similar phenotype was observed when wild-type FliM was expressed in the NMB155 cells. Wild-type FliM and FliM-E9K appear to be competitive in regard to determining flagellar number. We also observed that NMB155 cells swam almost without any directional changes. It is worth noting that a very small fraction of *fliM* deletion cells possessed flagella (Fig. 3 and Fig. S3).

**Fig. 3.**
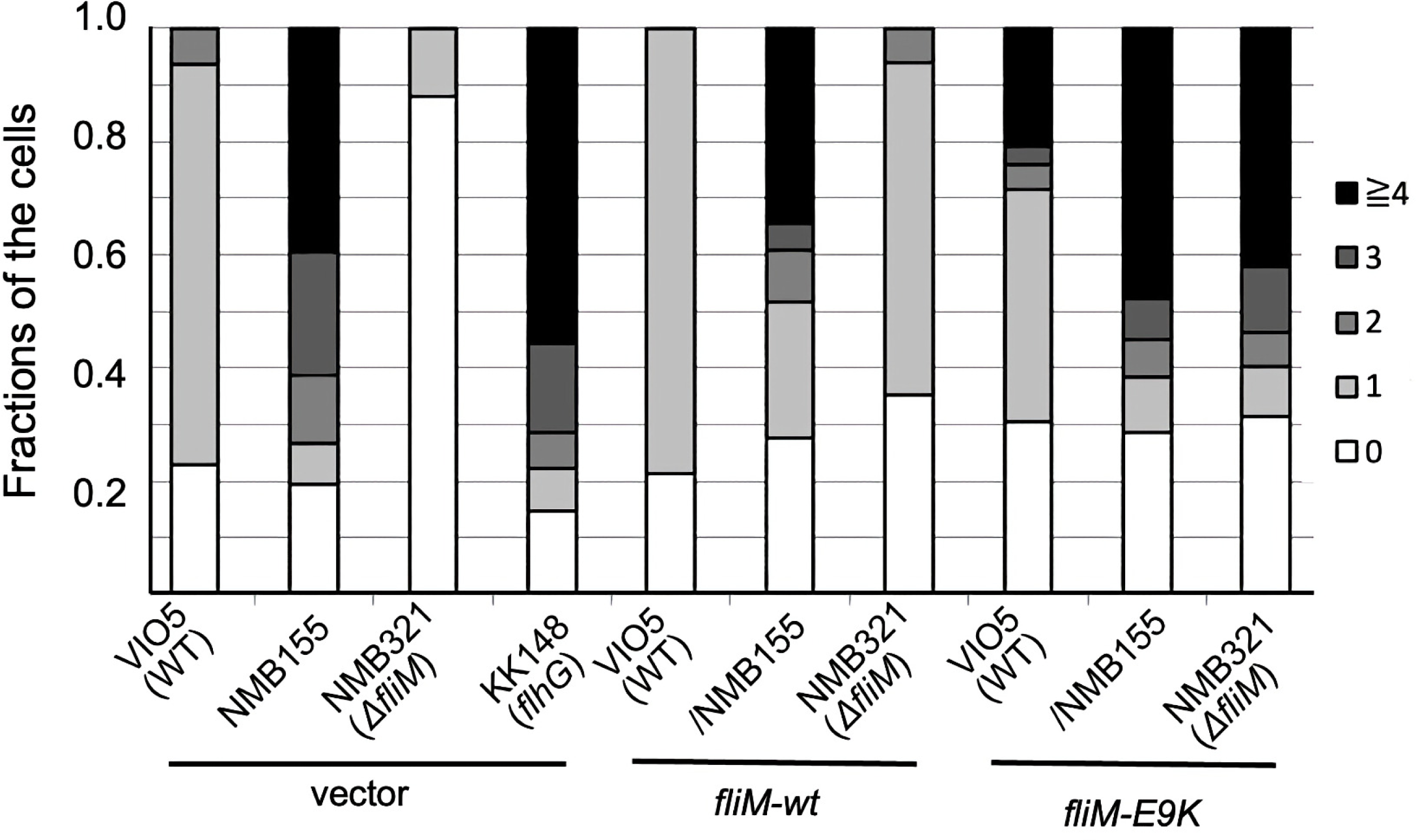
The flagellar number of cells. The cells for VIO5 (wild-type), NMB321 (Δ*fliM*), NMB155, and KK148 (*flhG*) containing pBAD33 (vector), pMK1001 (*fliM-wt*; expressing wild-type FliM), or pMK1001-E9K (*fliM-E9K*; expressing FliM-E9K) were cultured in VPG medium containing 0.02% (w/v) arabinose and 2.5 μg/mL of Cm at 30°C for 3 hours. The polar flagella were observed using a high-intensity dark-field microscope, and the number of flagella per every bacterial cell was measured and is presented as fractions of each population.

### The FliM-E9K mutation does not alter the polar localization of FlhG

Next, we examined the intracellular localization of FlhG. In wild-type cells, FlhG localizes to the cell pole and negatively controls flagellation. As the NMB155 and FliM-E9K mutant strains exhibited a multi-flagellar phenotype, we hypothesized that they reduced the polar localization of FlhG. To test this, the plasmid pAK541 encoding *flhG-gfp* was introduced into the wild-type (VIO5) and NMB155 and was observed under a fluorescence microscope. Fluorescence dots at the cell poles were observed in both wild-type and NMB155 cells expressing FlhG-GFP. We counted the number of individual cells in which fluorescent dots were observed and calculated the percentage of the total number of bacteria expressing FlhG-GFP (Fig. 4). Contrary to our hypothesis, we observed that the level of localization of FlhG-GFP in NMB155 was similar to that of VIO5, thus suggesting that FliM-E9K did not affect FlhG localization.

**Fig. 4.**
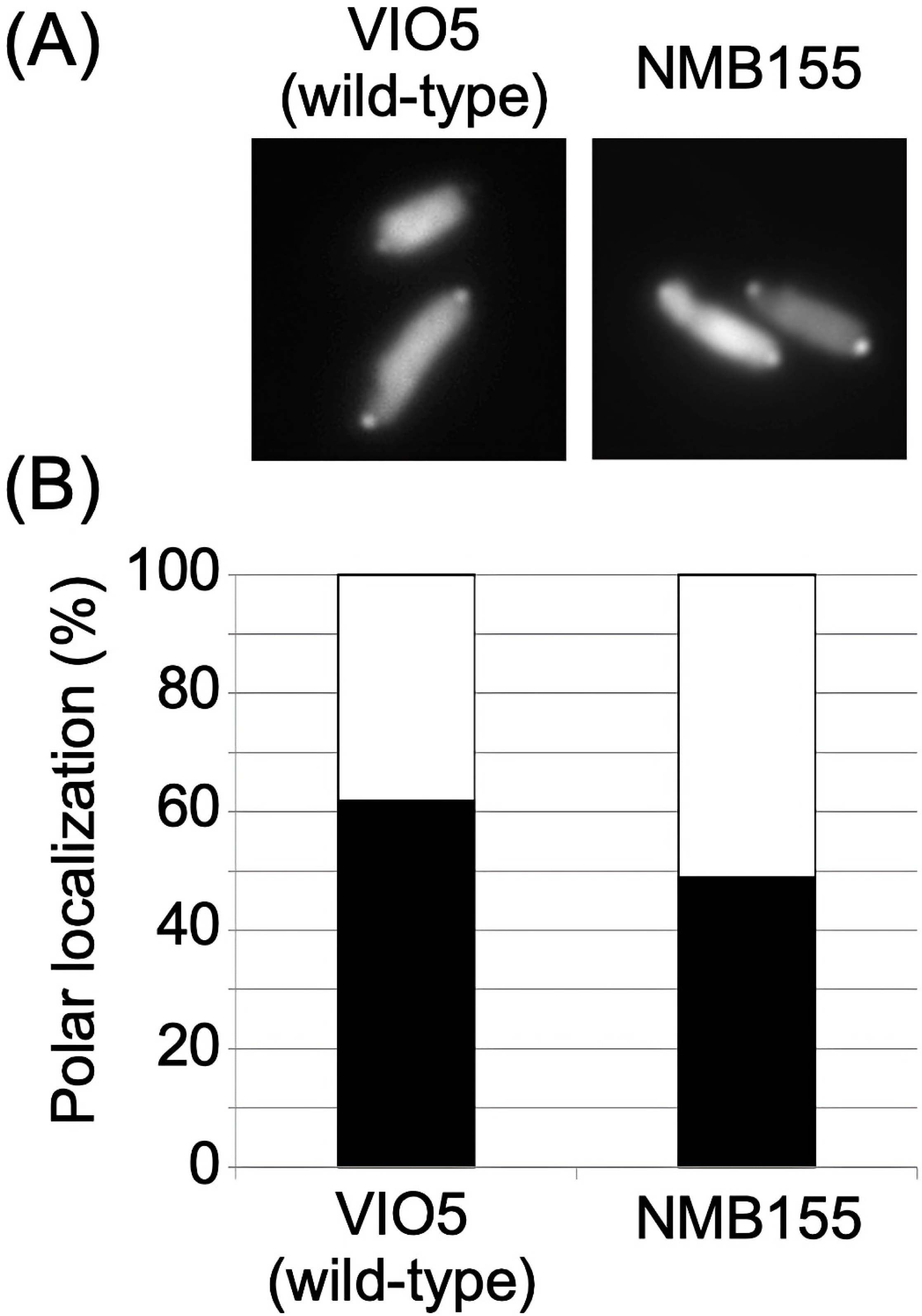
Observation of the polar localization of FlhG-GFP, VIO5 (wild-type), and NMB155 cells containing pAK541 encoding FlhG-GFP that were cultured in VPG medium containing 0.02% (w/v) arabinose and 2.5 μg/mL of Cm at 30°C for 4 hours. The samples were then subjected to a fluorescence microscopy (A) to count the number of the cells with fluorescent dots at the cell poles (B). The percentage of cells with fluorescent dots at the cell pole in the total population was provided.

### Mutation of the N-terminal CheY-P binding site of FliM

The E9 residue in FliM was highly conserved among FliMs from various bacterial species (Fig. 5A). Adjacent to the N-terminal side of E9, there is a residue that is also negatively charged (D8) but not as highly conserved. The crystal structure of the complex of the N-terminal peptide of FliM and CheY has been previously reported, and the residues corresponding to our E9 and D8 residues were oriented in the opposite direction from the contact surface with CheY (Fig. S4). Therefore, we speculated that the E9 and D8 residues may function cooperatively. To test this hypothesis, we generated and analyzed new mutations that included D8K, E9A, D8A, and E9A (Fig. 5B and 5C). D8K, where the charge of the Asp residue was reversed, exhibited the same swimming ability as did the wild type. This indicated that the FliM-D8 residue did not function like the FliM-E9 residue and that the FliM-E9 residue functions independently of the FliM-D8 residue to control flagellar synthesis. The E9A and D8A&E9A mutants conferred similar phenotypes with smaller motility rings than were observed with the wild-type and with larger expansion than was observed with E9K. High-intensity dark-field microscopy revealed that most cells treated with FliM-E9K possessed three or more flagella, and the E9A mutant also possessed multiple flagella that were similar to those of E9K. Under dark field microscopy, the E9K mutant cells exhibited few changes in swimming direction, while the E9A cells exhibited more frequent changes in swimming direction. The ability of the flagellar motor to control rotational direction explains the motility in the soft-agar plate of the FliM-E9A mutant.

**Fig. 5.**
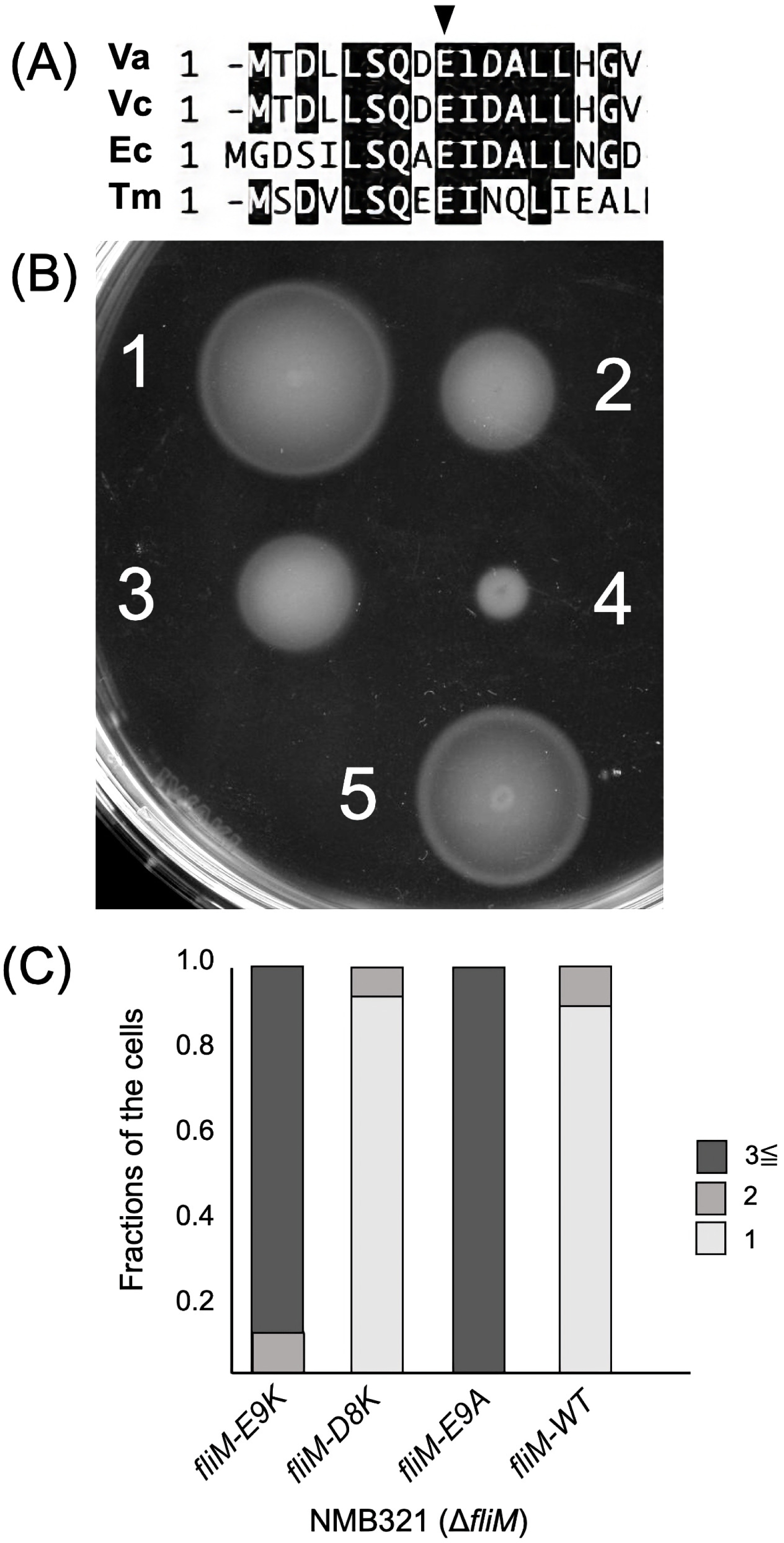
(A) Amino acid sequence alignment of the N-terminal region of FliM from four bacteria that include *V. alginolyticus* (Va), *V. cholerae* (Vc), *E. coli* (Ec), and *T. maritima* (Tm). The position of E9K in *V. alginolyticus* FliM is indicated by an arrowhead. (B) The motility of the FliM mutants. The fresh colonies of NMB321 (Δ*fliM*) containing plasmid expressing *fliM* (5; pMK1001) with or without *fliM* mutations (1; D8K, 2; E9A, 3; D8A&E9A or 4; E9K) were inoculated on VPG soft-agar plate containing 0.02% (w/v) arabinose and 2.5 μg/ml of Cm,and then incubated at 30°C for 6 hours. (C) The flagellar number of cells. The cells of NMB321 (Δ*fliM*) containing plasmid expressing *fliM* (pMK1001) with or without mutations (D8K, E9A or E9K) were cultured in VPG medium containing 0.02% (w/v) arabinose and 2.5 μg/ml of Cm at 30°C for 4 hours. The cells with polar flagella were counted using a high-intensity dark-field microscope, and the number of flagella per each flagellated cell was measured and is presented as fractions of each population.

These results indicate that FliM-E9 residues are important for both directional changes in flagellar motor rotation and the control of flagellar number. The charge-reversal E9K mutation affected both rotational direction control and flagellar number control, and the E9A mutation affected only flagellar number control and not rotational direction control.

### The properties of FliM-E9K protein

It is difficult to explain why the mutation in the rotator protein FliM confers the multi-flagellation phenotype. To test this hypothesis, we purified wild-type FliM and mutant FliM proteins and compared them. The FliM fragment lacking the N-terminal and C-terminal regions was cloned, and the crystal structure was resolved (27). However, the full length was not cloned, and therefore the *fliM* gene was cloned into the pET15b plasmid vector to express FliM with an N-terminal His-tag. After confirming expression and purification, the E9K mutation was introduced into this plasmid using a quikchange procedure. The wild-type FliM and FliM-E9K mutant proteins were purified using the His-tag, and the proteins were obtained. They were subjected to size-exclusion chromatography, and their elution profiles were identical (Fig. S5). Both of the proteins were separated as the main peak at approximately 90 kDa. As the molecular weight of His-tagged FliM is 41,979 based on estimations from the amino acid sequence, purified FliM appears to be present primarily as a dimer in solution. We detected small peaks at ca. 43 kDa and 350 kDa, thus suggesting that they corresponded to the monomer and multimer of FliM, respectively. We did not observe any differences between wild-type and mutant FliM-E9K proteins during the purification process. The structure of FlhG from *Geobacillus thermodenitrificans* has been resolved, and FlhG interacts with the FliM/FliY of C-ring proteins (23). Thus, we examined the binding of wild-type FliM and FliM-E9K to FlhG. Purified FlhG and wild-type FliM or FliM-E9K were mixed and analyzed using size-exclusion chromatography. No differences were detected based on the elution pattern (Fig. S5).

Next, the effects of wild-type FliM and FliM-E9K on the ATPase activity of FlhG were examined (Fig. 6). The ATPase activity of wild-type FlhG was low as reported previously (28) and released only a 5 µM concentration of phosphate (Fig. 6B). However, the addition of wild-type FliM (FliM-WT) resulted in a 4-fold increase in phosphate release, thus suggesting that the FliM protein activated the ATPase activity of FlhG (Fig. 6B, FliM-WT). In contrast, when the FliM-E9K mutant protein was mixed, no activation effect was observed (Fig. 6B, FliM-E9K). The D171A mutation in FlhG is known to enhance ATPase activity by up to 7-fold. The addition of FliM-WT or FliM-E9K did not further enhance ATPase activity in the FlhG-D171A mutant (Fig. 6C).

**Fig. 6.**
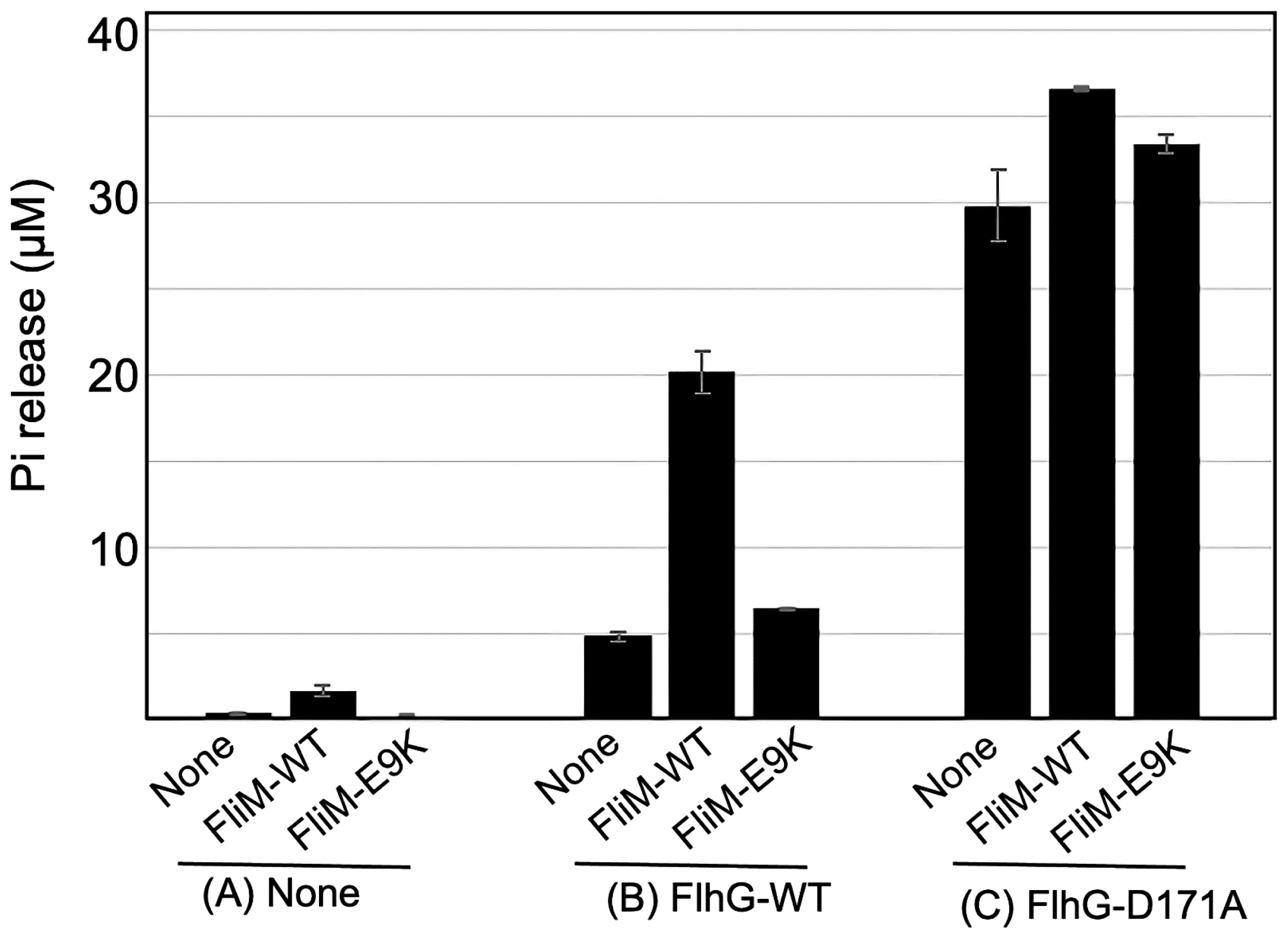
ATPase activity of FlhG. The buffer (None) or 2.5 μM FliM protein solutions (FliM-WT or FliM-E9K) were mixed with the buffer (A: None) or 5 μM FlhG protein solutions (B: FlhG-WT, or C: FlhG-D171A). Pi release was measured 30 min after the addition of 0.5 mM ATP. Activities were determined in 3 independent experiments. The error bars reveal the standard deviation of the mean value.

## Discussion

FlhF and FlhG are responsible for the essential regulation of flagella in *V. alginolyticus*. FlhF positively regulates flagellar formation, and FlhG negatively regulates FlhF function by reducing the number of flagella (19). FlhG is an ATPase and FlhF is a GTPase, an each may regulate the function of each other by influencing the activities of NTPase to reduce the number of flagella to one. Additionally, a polar-landmark protein HubP is required for the appropriate functions of FlhF and FlhG. The polar localization of FlhG is dependent upon HubP, and the HubP-deficient strain possessed multiple flagella in *V. alginolyticus* (25). Approximately 200 amino acids at the C-terminus of HubP are essential for its interaction with FlhG in *V. cholerae* (29). Furthermore, FlhG interacts with FliM, and N-terminal deficiency in FliM confers the phenotype of multiple flagella in *S. putrefaciens* (23, 24). These reports revealed that the master transcription factor FlrA (FlaK in *V. alginolyticus*) interacts with FlhG using the same motif as FliM. In *P. aeruginosa*, homologs of FlrA and FlhG (FleQ and FleN, respectively) have been studied in terms of transcriptional regulation of genes rather than the regulation of flagellar number (30, 31).

Among the cells isolated as mutants based on reduced motility on soft-agar plates, those with increased numbers of flagella, short-length flagella, and long-length flagella have been isolated for *V. alginolyticus* (26, 32). Mutations that cause abnormal flagellar length are not yet known. In this study, a novel mutation increases the number of flagella, and this mutation did not occur on *flhF* and *flhG* genes and was instead identified on the *fliM* gene. Specifically, the E9K mutation in FliM (FliM-E9K) increased the number of flagella. Initially, it was a complete mystery in regard to if *fliM* was the causative gene that increased the number of flagella. We hypothesized that the E9K mutation was responsible for promoting flagellar formation.

Flagellar formation begins with the formation of an export gate complex followed by MS-ring formation by FliF. FliM is thought to interact with FliG and FliN to form a C-ring after MS ring formation. The co-expression of FliG with FliF from *V. alginolyticus* promoted MS-ring formation in *E. coli* cells (33), thus suggesting that FliG promotes MS-ring formation or increases the stability of MS-ring in *V. alginolyticus* cells. Therefore, we speculated that the conformational change in the C-ring would affect the interaction between the MS-ring and C-ring and would promote the formation of multiple MS-ring. In contrast, the interaction of FlhG and FliM and the number of flagella were altered by mutations in the N-terminus of FliM in *B. subtilis* and *S. putrefaciens* (24). Mutation of FliM results in the formation of multiple flagella in *C. jejuni* (34, 35). The N-terminal region of FliM (FliM_N_) contains a highly conserved EIDAL motif that provides a high-affinity binding site for CheY-P. Deletion of the FliM_N_ region confers multiple flagellations, and FlhG does not localize to the cell pole due to a lack of the EIDAL motif in FliM in *S. putrefaciens* cells (23, 24). In *V. alginolyticus*, the FliM deletion strain exhibited no motility on the soft-agar plate; however, we detected a small fraction of motile cells in liquid medium and cells with flagella by observation using an electron microscope (Fig. S3). The FliM-E9K mutation occurred in the conserved EIDAL motif.

In this study, we demonstrated that FlhG was correctly localized at the cell pole in the FliM-E9K mutant of *V. alginolyticus*. Due to this mutation, chemotaxis was abnormal, thus suggesting that the binding of CheY-P to FliM was weakened. The ATPase activity of FlhG was increased by the addition of the FliM protein *in vitro* (Fig. 6). However, the direct interaction between purified FliM and FlhG was not detected according to size-exclusion chromatography (Fig. S5), thus indicating that the FliM-FlhG interaction was not strong. The ATPase activity of FlhG is important for the negative regulation of flagellar number, and the increase in ATPase activity resulting from a mutation (FlhG-D171A) strengthens the negative regulation by FlhG (28). Similarly, the FliM_N_ region appears to strengthen the negative regulation of flagellar formation by FlhG, as the FliM_N_ region increases the ATPase activity of FlhG. The FliM_N_ region is structurally flexible and is exposed in the outward direction of the C-ring. Therefore, the FliM_N_ region may prevent further formation of new flagella at the cell pole where the flagellum (or C-ring) has already formed. The FliM-E9K and E9A mutations may disturb the function of FliM. How the enhancement of FlhG ATPase activity by FliM is involved in flagellar number regulation remains unknown currently and will be addressed in future studies.

The crystal structure of FlhG of *Geobacillus thermodenitrificans* has been reported (23), the interaction with FliM was examined. The residues K177, R207, and F215 in the α6 and α7 helices in FlhG were demonstrated to be important for binding (Fig. S4C, S4D, Fig. S6). This FliM-binding region also competitively binds to the HTH domain in FlrA (FlrA-HTH) (24). Addition of FlrA-HTH increased FlhG ATPase activity. As *V. alginolyticus* FlhG is presumed to possess a similar structure to that of *G. thermodenitrificans* FlhG, it is likely that the FliM_N_ region directly interacts with the α6 and α7 helices of FlhG. The conformational change in the α6 and α7 helices of FlhG due to this interaction may increase the ATPase activity of FlhG, as the D171A mutation has been observed to increase ATPase activity in *V. alginolyticus* FlhG (28). As the D171 residue of *V. alginolyticus* FlhG is predicted to be located in the α6 helix, it is likely that conformational changes similar to those caused by the D171A mutation were induced by its interaction with the FliM_N_ region.

As mentioned above, it has been reported that the FliM_N_ region is highly conserved and functions as a CheY-P-binding site (36). In the report, the FliM-E10G of *Salmonella* that is mutated at residues corresponding to *V. alginolyticus* FliM-E9K or E9A and exhibits a marked reduction in CheY activity that is considerbly less than that of the wild type. The co-crystal structures of 16 residues peptide in FliM_N_ of *E. coli* FliM and CheY have been reported (37, 38), and the co-crystal structure of *Thermotoga maritima* CheY and FliM_N_ fragments has also yielded similar interaction results (39) (Fig. S4). The FliM_N_ region binds to CheY-P with high affinity, and CheY binds to FliN and the middle domain of FliM with low-affinity (40). Loss of the FliM_N_ region can also cause a CheY-mediated chemotaxis response based on the binding of CheY-P to the low-affinity sites and not to the high-affinity site, and this plays a major role in the orientation of the rotational direction of the flagellar motor (41). For flagellar formation, the FliM_N_ region is exposed outside of the C-ring structure. Thus, CheY-P binds and changes the rotational direction of the flagellar motor. The binding of CheY-P in proximity to the motor was demonstrated by GFP-fused CheY (42). *The E. coli* response regulator protein CheY is structurally similar to the Ras-like GTPase domain (43-45). The Ras-like GTPases and SIMIBI proteins belong to a superfamily of GTP-binding (G) proteins that constitute a class of P-loop (phosphate binding loop) proteins that function as molecular switches between the NDP-bound state and the NTP-bound state (46, 47). FlhG belongs to the same superfamily. The N-terminal fragment of FliM is involved in the regulation of P-loop proteins to switch between the active and inactive states in the superfamily.

## Materials and Methods

### Bacteria strain and plasmid

Bacterial strains and plasmids are listed in Table S1. *E. coli* was cultured in LB medium (1% [w/v] bactotryptone, 0.5% [w/v] yeast extract, and 0.5% [w/v] NaCl]). *V. alginolyticus* was cultured at 30°C in VC medium (0.5% [w/v] hipolypeptone, 0.5% [w/v] yeast extract, 0.4% [w/v] K_2_HPO_4_, 3% [w/v] NaCl, and 0.2% [w/v] glucose) or VPG medium (1% [w/v] hipolypeptone, 0.4% [w/v] K_2_HPO_4_, 3% [w/v] NaCl, and 0.5% [w/v] glycerol). Chloramphenicol (Cm) was added to a final concentration of 25 µg/mL, and 2.5 µg/mL was used for *E. coli* and *V. alginolyticus*, respectively. Ampicillin (Amp) was added at a final concentration of 100 µg/mL to *E. coli*.

### Mutagenesis

Site-directed mutagenesis was performed using the QuikChange site-directed mutagenesis method as described by Agilent Technologies (Santa Clara, USA). Transformation of *V. alginolyticus* using pMK1001 and mutant derivative plasmids was performed using electroporation (48).

### Swimming assay in the soft-agar plate

*V. alginolyticus* cells were plated on VPG agar plates (VPG containing 1.25 % [w/v] agar) with antibiotics. A colony of *V. alginolyticus* was inoculated onto VPG soft-agar plates (VPG containing 0.3% [w/v] bactoagar and 0.02% [w/v] arabinose) and incubated at 30°C for 4 h.

### Measurement of the number of polar flagella

*V. alginolyticus* cells were cultured overnight in VC medium, diluted 50-fold in VPG medium containing 0.02% (w/v) arabinose, and incubated at 30°C for 3 h. Cells (1 mL) were collected by centrifugation at 3,500 × *g* for 5 min at 4°C. The cells were washed with 1 mL of TMN50 (50 mM Tris-HCl [pH 7.5], 5 mM glucose, 5 mM MgCl_2_, 50 mM NaCl, 250 mM KCl), and then suspended in 500 μL of TMN50. This cell suspension was diluted 5 times with TMN50, and CCCP was added to a final concentration of 200 μM. Each prepared sample (6 μL) was spotted on a glass slide, covered with a cover glass, the cells were observed using a high-intensity dark-field microscope (Olympus model BHT), and the number of polar flagella was counted. The number of flagella observed was classified into five categories that included 0, 1, 2, 3, and ≥ 4. The percentages were calculated by adding the results of three independent experiments where greater than 50 individual cells were observed per sample.

### Fluorescence microscopy

*V. alginolyticus* cells cultured overnight in VC medium were diluted 50-fold in VPG medium containing 0.02% (w/v) arabinose and were incubated at 30°C for 4 h. Cells (1 mL) were collected by centrifugation at 3,500 × *g* for 5 min, suspended in 200 μL of V buffer (50 mM Tris-HCl [pH8.0], 5 mM MgCl_2_, 300 mM NaCl), and used as samples. A gap was made between the glass slide and the cover glass using double-sided tape. The gap was filled with 0.1% (w/v) poly-l-lysine and allowed to stand for 1 min. Subsequently, 20 μL of V buffer was perfused to wash the gap. Next, 20 μL of the sample was poured into the gap and allowed to stand for 1 min to fix the cells, and 20 μL of V buffer was perfused to remove the unattached cells. The samples were observed under a fluorescence microscope (Olympus BX50). The polar localization rate of FlhG was defined as the ratio of the number of individuals in which the fluorescent dots were visible only at the cell poles to the total number of individuals.

### Purification of FlhG protein

*E. coli* BL21(DE3)/pLysS was transformed with pTrc-flhG (encoding wild-type FlhG) or its derivative (encoding mutant FlhG-D171A). Proteins were purified as described previously (49).

### Purification of FliM protein

*E. coli* BL21(DE3) was transformed with pMK2001 (encoding wild-type FliM) or its derivative (encoding mutant FliM-E9K). Cells were cultured in LB medium containing ampicillin at 37°C until 0.8 to 1.0 at OD 660 nm was achieved. Then, the culture was cooled in iced water for 20 min, isopropyl-β-_D_-thiogalactopyranoside (IPTG) was added to a final concentration of 0.4 mM, and the culture was incubated at 16°C overnight. The cells were collected by centrifugation at 7,000 × *g* for 10 min at 4°C and suspended in 20 mL of 20TN150 (20 mM Tris-HCl (pH8.0), 150 mM NaCl) containing 1 mM ethylenediaminetetraacetic acid (EDTA). The cells were then sonicated (large probe, power=8, 60 s, 3 times, duty cycle 50%). The unbroken cells were removed by centrifugation at 16,000 × *g* for 10 min at 4°C, and the cells were resuspended in 20 mL of 20TN150 containing 1 mM EDTA, where sonication was repeated. The resulting supernatants were collected, MgCl_2_ and imidazole were added at a final concentration of 2 mM and 10 mM, respectively, they were then centrifuged at 13,500 × *g* for 10 min at 4°C, and the supernatant was further ultracentrifuged at 154,000 × *g* for 60 min at 4°C. The soluble fraction of this sample was loaded onto a HiTrap TALON column (5 mL, GE Healthcare) connected to an ÄKTAprime system (GE Healthcare) and washed with 20TN150 buffer containing 5 mM MgCl_2_ and 20 mM imidazole. The proteins were eluted and collected using an imidazole gradient. The peak fractions were collected, frozen in liquid nitrogen, and stored at -80°C.

### Size-exclusion chromatography

The mixture of purified FliM solution and the purified FlhG solution or the buffer was ultracentrifuged to remove aggregated materials, and the 100 μL volume from the supernatant was injected into the size exclusion chromatography (Superdex 200 10/300) and eluted by 0.5 mL/min with 20TN150 buffer containing 5 mM MgCl_2_ and 10% glycerol.

### Measurement of ATPase activity of purified FlhG

The ATPase activity of FlhG proteins was evaluated by measuring the concentration of inorganic phosphate produced by the degradation of ATP when mixed with ATP using a kit from Innova Bioscience as previously described (28).

## Acknowledgements

We thank Dr. Kimika Maki for technical support with electron microscopy. This work was supported in part by JSPS KAKENHI Grant Numbers 16H04774 (to S.K.) and 20H03220 (to M.H.).

## Supporting information

The supplementary information associated with this article can be located in the online version of the publisher website.

## Figure legends

**Fig. S1.** Schematic illustration of the polar flagellar motor of *V. alginolyticus*. (A) The genomic context of the flagellar genes around *fliM*. (B) The vertical section image of the flagellar motor. The C-ring is composed of FliG, FliM and FliN, the MS-ring is composed of FliF, and the stator is composed of PomA and PomB. FliM is indicated by red.

**Fig. S2.** (A) The model of the C-ring. Atomic models of FliG (green), FliM (blue), CheY (yellow), and C-terminal region of FliF (light purple) were fitted into the electron microscopy map of C-ring. The structure was referred to for our previous paper (Nishikino *et al*., 2018, *Sci Rep* 8:17793). (B) Amino acid sequence alignment of FliM from four bacteria, including *V. alginolyticus* (Va), *V. cholerae* (Vc), *E. coli* (Ec), and *T. maritima* (Tm) is provided. The identical and homologous residues were highlight in black and gray boxes, respectively. The position of E9 in *V. alginolyticus* FliM was indicated by an arrowhead.

**Fig. S3.** NMB321 (Δ*fliM*) was grown on VC hard-agar plate and cells were observed by electron microscopy.

**Fig. S4.** Atomic models of the CheY-FliM_N_ complex and FlhG. (A, B) The ribbon model of the structure of complex of CheY and the N-terminal 16 residues peptide of FliM (PDB ID: 1U8T). The CheY is presented in yellow and FliM is presented in blue. The red filled spheres model is the corresponding residue to E9 of *V. alginolyticus* FliM. (A) and (B) were views from different angles. (C, D) The ribbon model of the structure of *G. thermodenitrificans* FlhG (PDB ID: 4RZ2). The FlhG is presented in yellow and the important residues for FliM-binding are presented in red. (C) and (D) were views from different angles.

**Fig. S5.** Size-exclusion chromatography. FliM only (A), the mixture of FliM and FlhG (B), the mixture of FliM, FlhG and ATP (C), or FliM-E9K only (D) were ultracentrifuged to remove aggregated materials, and the 100 μL volume from the supernatants were injected into the size-exclusion chromatography (Superdex 200 10/300). The black filled arrow, open arrow, and gray filled arrow indicate the peak positions whose estimated sizes were ca. 90 kDa, ca. 350 kDa, and ca. 26 kDa, respectively.

**Fig. S6.** Amino acid sequence alignment of FlhG homologs. (1) MinD (*E. coli*); (2) FlhG (*C. jejuni*); (3) FleN (*P. aeruginosa*); (4) FlhG (*V. alginolyticus*); (5) FlhG (*S. putrefaciens*); (6) FlhG (*G. thermodenitrificans*); (7) YlxH (*B. subtilis*). The interaction sites with FliM in *G. thermodenitrificans* are presented by open arrow heads. The position of D171 in *V. alginolyticus* FlhG is presented by a black filled arrowhead. The lines of red, blue, light blue, and purple on the sequences are regions for P-loop, Switch I, Switch II, and a C-terminal amphipathic helix, respectively

